# Dynamic histone hyperacetylation shapes environmentally responsive chromatin states

**DOI:** 10.64898/2026.06.19.732295

**Authors:** Moonia Ammari, Linkan Dash, Aanchal Choudhary, Hazel Mamania, Jeevika Gupta, Moorthy Gnanarajah, Kristoffer Gittens, Mark Zander

## Abstract

Transcription factors (TFs) orchestrate environmental responses by activating target genes, yet how they reshape epigenome architecture to coordinate gene expression remains poorly understood. We previously identified SIENA (Stimulus-Induced ENhancer Acetylation) domains as large regions of jasmonic acid (JA)-induced H3K9 hyperacetylation surrounding MYC2 TF binding sites in *Arabidopsis* and tomato. However, the mechanisms underlying the formation of SIENA domains (SIENAs) and their functional significance remained unknown. Here, we show that SIENAs also form at major JA-responsive genes and gene clusters in soybean, extending this phenomenon to an evolutionarily distant crop species. Comprehensive chromatin profiling revealed that SIENAs accumulate multiple histone acetylation marks, including H3K9ac, H3K27ac, H3K56ac, H2BK20ac, and H2A.Zac, establishing them as regions of broad histone hyperacetylation. Pharmacological disruption of proteasomal turnover and histone acetylation dynamics compromised SIENA formation. Chromatin accessibility analyses further showed that inducible accessibility within SIENAs is tightly associated with MYC2 binding sites, supporting a model in which MYCs nucleate localized chromatin reprogramming events. Together, our findings establish SIENAs as MYC2-dependent chromatin-organizing domains and identify histone hyperacetylation as a central feature of MYC2-mediated gene activation.

## INTRODUCTION

Plants continuously adapt their growth, development, and defense programs to changing environmental conditions. These adaptive responses require rapid and coordinated transcriptional reprogramming that enables plants to dynamically respond to biotic and abiotic stimuli (Tsuda and Somssich, 2015, Hickman et al., 2017, Song et al., 2016). TFs play a central role in this environmental responsiveness by serving as molecular platforms assembling a diverse array of chromatin regulators (CRs), that modulate the transcriptional output of TF-target genes via epigenome reprogramming (Ammari et al., 2024). Features of the epigenome that are dynamically and reversibly regulated by the concerted interplay of TFs and CRs are genome-wide occupancies of histone tail post-translational modifications (PTMs) and histone variants, DNA methylation, chromatin accessibility, and three-dimensional (3D) chromatin architecture (Lloyd and Lister, 2022). The acetylation of various lysines at histone tails represents one of the most prominent histone PTMs and a hallmark of transcriptionally permissive chromatin. Histone H3 and H4 tail acetylation can weaken the electrostatic interactions between histones and nucleosomal DNA which in turn leads to increased chromatin accessibility for the transcriptional machinery (Chen et al., 2024). The dynamic balance of histone acetylation is established and maintained through the antagonistic activities of histone acetyltransferases (HATs) and histone deacetylases (HDACs), which function within multi-protein complexes capable of targeting multiple lysine residues across histone tails (Chen et al., 2024).

The JA signaling pathway represents an excellent model for investigating environmentally responsive histone acetylation dynamics in plants. As a central phytohormone regulating plant immunity, JA orchestrates extensive defense-associated gene regulatory networks (GRNs), which is accompanied by widespread reprogramming of the histone acetylation landscape (Zander et al., 2020, Choudhary et al., 2024, Hickman et al., 2017). JA is perceived in its bioactive conjugated form, JA-Ile ((+)-7-iso-jasmonoyl-L-isoleucine), directly on chromatin by a co-receptor complex consisting of the F-box protein COI1 (CORONATINE INSENSITIVE 1), a component of the SKP1-Cullin F-box E3 ubiquitin ligase complex (SCF^COI1^), and JAZ (JASMONATE ZIM-DOMAIN) repressor proteins (JAZs) (Thines et al., 2007, Chini et al., 2007, An et al., 2017). JA-Ile perception by the SCF^COI1^-JAZ co-receptor complex centers around the master TF MYC2 (MYELOCYTOMATOSIS 2) and its closely related homologs MYC3, MYC4, and MYC5 (MYCs) (Qi et al., 2015, Lorenzo et al., 2004, Fernandez-Calvo et al., 2011, Song et al., 2017), ultimately determining the regulatory complexes with which MYCs associate. Under low nuclear JA-Ile levels, the transcriptional activator function of MYCs is repressed by JAZs through their association with the TPL (TOPLESS) co-repressor complex via NINJA (NOVEL INTERACTOR OF JAZ) (Pauwels et al., 2010). Upon increasing JA-Ile levels, JAZs become ubiquitinated and subsequently degraded by the 26S proteasome, thereby releasing MYCs to associate with the permissive MED (Mediator) complex containing MED16, MED25, HAC1 (HISTONE ACETYLTRANSFERASE OF THE CBP FAMILY1) and the Gro/Tup1 family proteins LUG (LEUNIG) and LUH (LEUNIG_HOMOLOG) (An et al., 2017, Chen et al., 2012, You et al., 2019, Wang et al., 2019).

Intriguingly, one of the major chromatin features regulated by the restrictive TPL and permissive MED complexes in a JA-dependent manner are histone acetylation dynamics (Zander and Vesper, 2026). Activation of JA signaling is accompanied by increased histone acetylation across JA-induced genes, whereas JA-mediated transcriptional repression correlates with corresponding decreases in histone acetylation levels (Choudhary et al., 2024). Notably, these chromatin changes are highly dynamic and rapidly return to basal levels following JA withdrawal (Choudhary et al., 2024). Despite the extensive nature of this JA-induced chromatin reprogramming, the underlying molecular mechanisms are only partially understood. The central role of MYC TFs in this process becomes evident from the observation that JA-induced H3K9 acetylation is largely abolished in *Arabidopsis myc2 myc3 myc4* (*myc234*) triple mutants (Choudhary et al., 2024), emphasizing MYCs’ role as a critical regulatory hub linking JA perception to epigenome reprogramming.

To date, only two HATs, HAC1 and GCN5 (GENERAL CONTROL NONDEREPRESSIBLE 5), have been implicated in JA signaling (Liu et al., 2019, An et al., 2017, An et al., 2022). HAC1 is part of the MED25-HAC1-LUH module within the permissive MED complex, linking JA-Ile perception-mediated liberation of MYCs to the reprogramming of the histone acetylation landscape (Liu et al., 2019). However, *hac1* mutants display relatively mild defects in JA-induced H3K9 acetylation (Choudhary et al., 2024). In contrast, GCN5 negatively regulates JA signaling through direct acetylation of TPL, thereby enhancing TPL-mediated transcriptional repression (An et al., 2022). On the other hand, the two reported histone deacetylases (HDACs), HDA6 (HISTONE DEACETYLASE 6) and HDA19, positively regulate JA signaling by deacetylating TPL, which promotes dissociation of the TPL complex from JAZ repressors (An et al., 2022). In addition, the simultaneous mutations of *TPL*, *TPR1* (*TOPLESS-RELATED 1*), and *TPR4*, mutation of *NINJA*, as well as higher-order mutations affecting ten *JAZ* genes, results in accelerated JA-induced H3K9ac accumulation, supporting their roles as negative regulators of histone acetylation (Choudhary et al., 2024). Nevertheless, despite the reported functions of HAC1, GCN5, HDA6, and HDA19, the minor phenotypes strongly indicate the involvement of additional operational HATs and HDACs during active JA signaling.

Importantly, the genomic location of histone acetylation is critical for its potential regulatory function. Histone acetylation can occur within gene bodies - typically showing the strongest enrichment around the +1 nucleosome- or within distal regulatory and enhancer regions (Chen et al., 2024). Most reports describing dynamic JA-responsive histone acetylation have focused on gene bodies, making it difficult to disentangle the direct regulatory contribution of histone acetylation from transcription-associated effects. Previously, we identified JA-induced domains of H3K9 hyperacetylation in regulatory regions of JA-responsive genes in both *Arabidopsis* and tomato (*Solanum lycopersicum*) which we termed SIENAs (Choudhary et al., 2024). These SIENAs can span entire defense-associated gene clusters in tomato and occur around MYC2-associated *cis*-regulatory elements (CREs) (Choudhary et al., 2024). Their absence in *Arabidopsis* JA-treated *myc234* mutants suggests that DNA-bound MYCs are essential for their formation (Choudhary et al., 2024).

Here, we investigate the mechanistic and structural role of SIENAs during active JA signaling. Through comparative analyses of H3K9ac landscapes in *Arabidopsis*, tomato, and soybean (*Glycine max*), we identify SIENAs as a conserved feature of a subset of MYC2 target genes. SIENAs can occur at individual genes or span entire gene clusters, thereby facilitating transcriptional co-regulation. By profiling multiple histone acetylation marks and chromatin accessibility, we show that SIENAs represent broad multi-lysine acetylation domains established through MYC-dependent chromatin reprogramming. Pharmacological perturbation of the JA pathway further reveals that proteasome-mediated degradation of JA signaling components is essential for SIENA formation, whereas inhibition of histone deacetylase activity supports a role for non-histone acetylation of TPL/TPR complexes in regulating these domains. Together, our findings establish SIENAs as conserved chromatin features that couple hormone perception to large-scale chromatin reprogramming and coordinated transcriptional activation.

## RESULTS

### SIENAs are conserved features of the JA-responsive chromatin landscape

SIENAs are characterized by JA-induced H3K9 hyperacetylation surrounding MYC2 binding sites within regulatory regions of JA-responsive gene clusters in *Arabidopsis* and tomato (Choudhary et al., 2024). To investigate whether SIENA domain formation extends beyond these species, we analyzed JA-induced H3K9ac dynamics in soybean, a phylogenetically distant legume crop species. *Arabidopsis*, tomato, and soybean seedlings were either left untreated or treated with JA for 2 h prior to H3K9ac profiling using PHILO ChIP-seq (Fig. 1A), and genes associated with JA-inducible H3K9ac domains were subsequently identified. The H3K9ac landscape was highly JA-responsive in all three species showing increased H3K9ac occupancy in their gene bodies after JA exposure (Fig. 1B and Supplementary Table S1).

**Figure 1.**
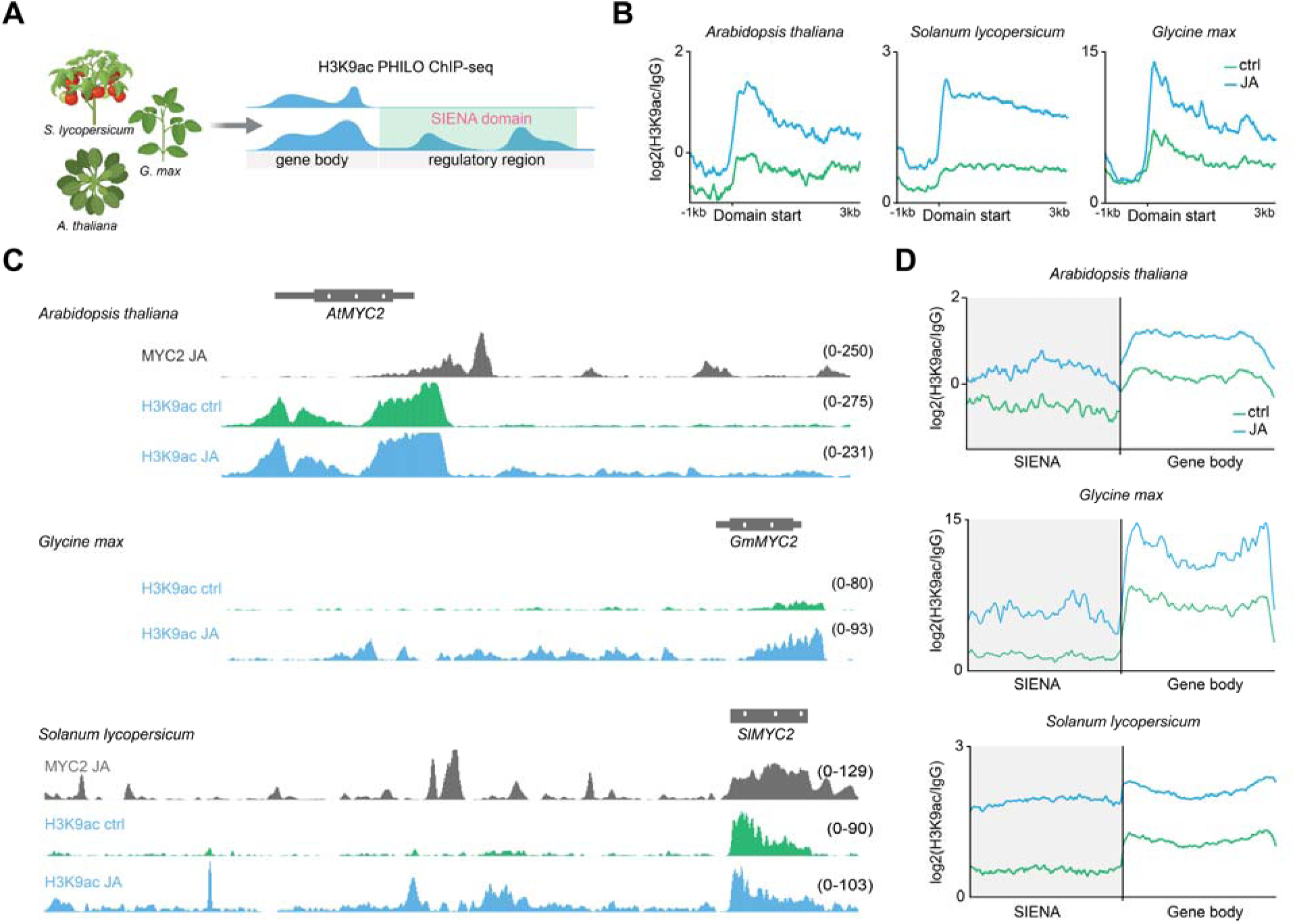
SIENAs are conserved features of the JA-responsive chromatin landscape. **A**, Schematic illustration shows experimental design for SIENA detection in the indicated plant species. Illustration was generated with BioRender (https://BioRender.com/9hf0evq). **B,** Metagene plots show JA-induced H3K9ac dynamics in *Arabidopsis thaliana*, *Solanum lycopersicum* and *Glycine max*. Plots are derived from two merged replicates and regions 1 kb upstream to 3 kb downstream of the domain start sites are shown. Enrichment levels were calculated as the ratio between merged H3K9ac samples and IgG control, and genomic location of plotted domains can be found in Supplementary Table S1. **C,** Genome browser screenshots showing SIENAs at the *MYC2* locus in *Arabidopsis*, tomato, and soybean. Each track represents the merged signal from two biological replicates and was normalized to sequencing depth. For *Arabidopsis* and tomato, MYC2 occupancy profiles are also shown. *Arabidopsis* MYC2 ChIP-seq data was obtained from Choudhary et al. (2024), and tomato MYC2 ChIP-seq data were obtained from Du et al. (2017). All tracks were normalized to their sequencing depth. **D,** Metagene plots show H3K9ac enrichment across SIENAs and associated gene bodies in untreated and 2 h JA-treated samples. Signals represent merged data from two biological replicates. SIENAs and gene bodies were scaled to 2 kb, and enrichment values were calculated relative to the corresponding IgG control.

We next investigated whether genes displaying JA-induced H3K9ac accumulation in their gene bodies also formed SIENAs which were defined as JA-inducible H3K9ac domains extending from gene bodies into regulatory regions, with a minimum size threshold of 1000 bp. Structurally, we distinguish between single-gene SIENAs and multi-gene SIENAs. Single-gene SIENAs, such as those associated with the *MYC2* gene (Fig. 1C), typically formed continuous H3K9ac domains extending from the gene body into the regulatory regions and, in some cases, even spreading into neighboring genes (Supplementary Fig. 1A, B). In contrast, multi-gene SIENAs extended across several genes, including both their regulatory regions and gene bodies, thereby forming large acetylated chromatin domains (Supplementary Fig. 1A, C). In *Arabidopsis*, we identified 30 SIENAs, compared with 114 in soybean and more than 800 in tomato, demonstrating extensive JA-induced reprogramming of the chromatin landscape in regulatory regions of responsive genes (Fig. 1C, D, Supplementary Fig. 1D and Supplementary Table S2). Using the size of JA-induced H3K9ac domains as a proxy for SIENA domain formation, we found that SIENAs in soybean were generally larger than those in *Arabidopsis* but remained considerably smaller than those observed in tomato (Supplementary Fig. 1D). Together, these analyses revealed substantial interspecies differences in both the abundance and size of SIENAs.

In *Arabidopsis*, which possesses a relatively compact genome, most SIENAs were restricted to single genes, with only two spanning multiple loci (Supplementary Fig. 1E). By contrast, tomato displayed substantially larger SIENAs, with some gene clusters encompassing up to seven genes within a single continuous acetylated domain (Supplementary Fig. 1E). In soybean, the SIENA structure resembled that of *Arabidopsis* with most SIENAs predominantly associated with single genes and only a few with multiple genes (Supplementary Fig. 1E). However, their sizes were considerably larger than those observed in *Arabidopsis* (Supplementary Fig. 1D).

As observed in *Arabidopsis*, where SIENAs occur at key MYC2 target genes within the core JA signaling network, gene ontology analysis of SIENA-associated genes in tomato and soybean, also revealed a strong enrichment for the JA signaling pathway (Supplementary Fig. 2A-C). A striking example of a gene associated with a substantial SIENA domain is *MYC2* itself, which is also one of the major targets of MYC2 (Du et al., 2017, Wang et al., 2019, Zander et al., 2020). Elucidating the chromatin architecture underlying this positive-feedback loop is particularly important given the central position of MYC2 at the apex of the JA GRN. The *Arabidopsis MYC2* regulatory region is unusually large, spanning approximately eight kilobases (kb) and containing multiple MYC2 binding sites, with the SIENA domain extending across the entire region (Fig. 1C). Notably, the *MYC2* regulatory regions in tomato and soybean also form prominent SIENAs with the tomato region exceeding 23 kb (Fig. 1C). Moreover, *JAZ* genes were associated with prominent SIENAs (Supplementary Fig. 2D). Among these, the soybean *GmJAZ10* locus displayed a ~17 kb SIENA domain, making it one of the most extensive SIENAs detected in soybean (Supplementary Fig. 2D and Supplementary Table S2). The recurrent formation of large SIENAs at *MYC2* and *JAZ* loci highlights a conserved chromatin architecture centered on the core regulatory circuitry of the JA pathway.

### Uniform formation of multi-gene SIENAs correlates with transcriptional co-regulation

To investigate the dynamic onset of JA-induced H3K9 acetylation at the existing SIENA genes and whether the H3K9ac increase in gene bodies and regulatory regions are temporally distinct processes, we profiled H3K9ac occupancy across a JA time series in tomato (control, 15 min, 30 min, 1 h, 2 h, and 2 h JA + 2h withdrawal (WD)) (Fig. 2A). Tomato was selected because it exhibits substantially larger SIENAs than *Arabidopsis*, characterized by large multi-gene SIENAs, and therefore provides an efficient system to capture coordinated acetylation dynamics across multiple regulatory regions and gene bodies (Supplementary Fig. 1D, E). We used two adjacent protease inhibitor (PI) clusters (PI I and PI II) as representative multi-gene SIENAs to illustrate both the large size of these domains and their dynamic formation following JA treatment (Fig. 2A). The most pronounced change occurred between 30 min and 1 h of JA treatment (Fig. 2A). Whereas little to no SIENA domain formation was detectable after 30 min of JA, SIENAs had already reached near-maximal levels by 1 h of JA (Fig. 2A). The substantial reduction in SIENA size following the 2 h JA withdrawal phase further highlights the highly dynamic and reversible nature of SIENA formation (Fig. 2A).

**Figure 2.**
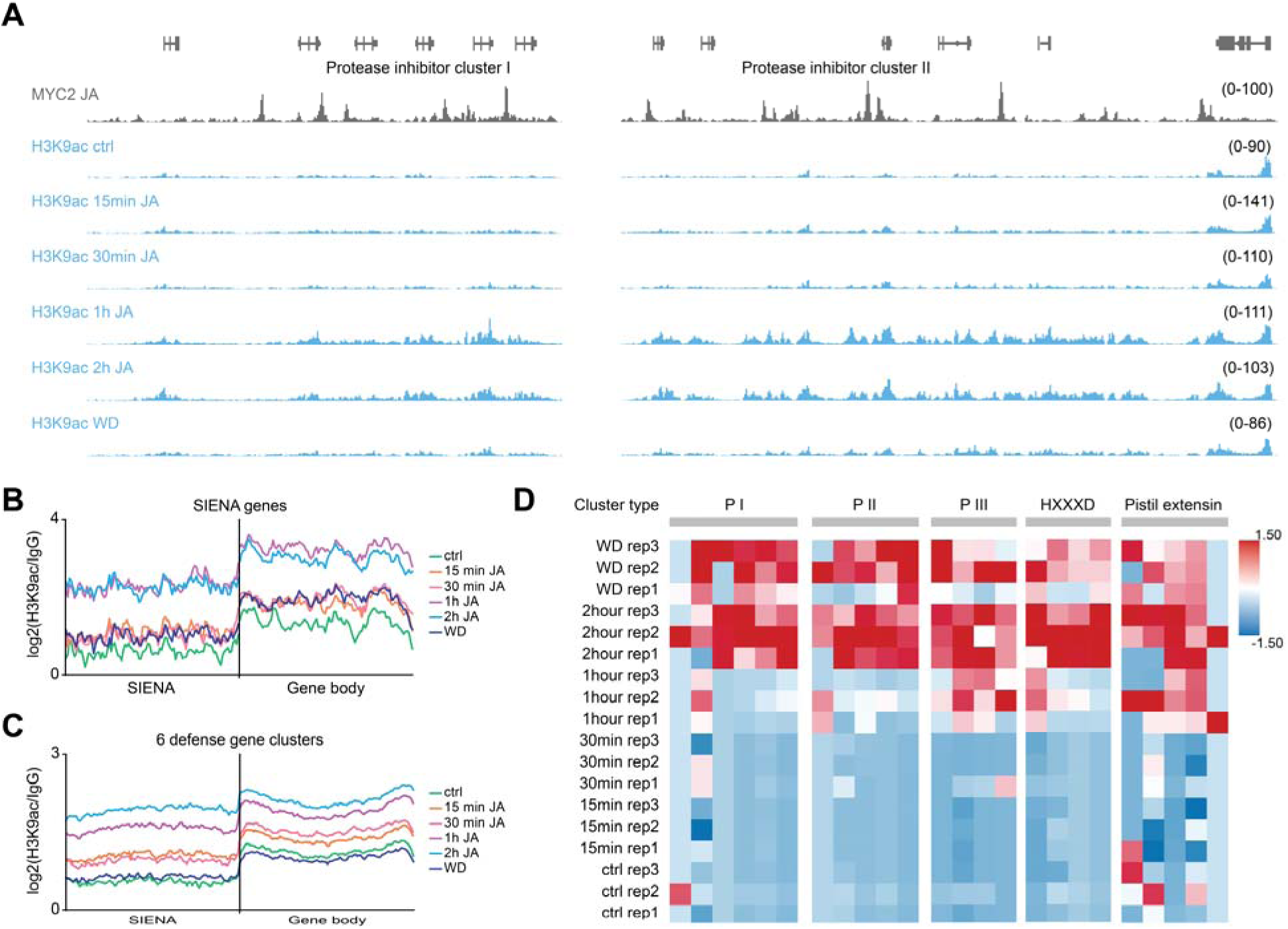
Uniform formation of multi-gene SIENAs correlates with transcriptional co-regulation. **A,** Genome browser screenshot illustrates H3K9ac dynamics across a JA time series (ctrl, 15 min, 30 min, 1 h, 2 h, 2 h + 2 h JA withdrawal (WD)) at two adjacent PI gene clusters (PI I and PI II). MYC2 occupancy profiles are included for comparison and were derived from the ChIP-seq dataset reported by Du et al. (2017). Each track represents the merged signal from two biological replicates and was normalized to sequencing depth. **B, C,** Metagene plots show the temporal dynamics of JA-induced H3K9ac enrichment across all tomato SIENA genes (**B**) and, separately, across genes located within the five selected multi-gene SIENAs (**C**). H3K9ac enrichment was calculated as the ratio of H3K9ac signal to the corresponding IgG control. For visualization, both gene bodies and SIENA regions were scaled to 2 kb. **D,** Heatmap shows expression dynamics for genes residing within multi-gene SIENAs determined with RNA-seq. Columns represent the JA time series (ctrl, 15 min, 30 min, 1 h, 2 h and WD), each in biological triplicate (rep1–rep3). Rows represent individual genes and colors denote row-scaled (*z*-score) normalized expression, from low (blue) to high (red). Genes are ordered and color-coded by cluster type (left): P I, P II, P III, HXXXD and pistil extension.

At the genome-wide level, analysis of 818 tomato SIENA-associated genes similarly revealed extensive JA-induced reprogramming of the H3K9ac landscape (Fig. 2B). We observed increased H3K9ac accumulation after 1 h of JA treatment which reached its maximum after 2 h, demonstrating the rapid establishment of SIENAs in response to JA (Fig. A, B). Increases in H3K9ac occurred concurrently in regulatory regions and gene bodies, although overall enrichment levels were generally lower in regulatory regions (Fig. 2A, B). To further investigate these dynamics in multi-gene SIENAs, we focused on five large defense-related gene clusters showing highly JA-inducible SIENAs, including three protease inhibitor clusters (PI I-III), a HXXXD-type acyltransferase cluster, and a pistil extension cluster (Supplementary Table S3). Across all five clusters, the first robust increase in H3K9ac was also observed after 1h JA and was not further enhanced in a 2 h JA period (Fig. 2A, C). The JA-induced increase of H3K9ac levels occurred also simultaneously within SIENAs and their corresponding gene bodies, suggesting the existence of a shared regulatory mechanism coordinating histone acetylation in gene bodies and corresponding regulatory regions.

Next, we investigated the transcriptional consequences of SIENA formation for genes residing within multi-gene SIENA domains in our tomato JA time-series experiment. Focusing on the five defense-associated gene clusters that displayed substantial SIENA domain formation, we found that genes embedded within these clusters were uniformly induced by JA with transcript levels generally peaking 2 h after JA treatment displaying a remarkable degree of transcriptional co-regulation (Fig. 2D). Notably, the onset of transcriptional induction was slightly delayed relative to the accumulation of H3K9ac. Whereas robust JA-induced gene expression was first detected after 2 h in four of the five clusters examined, H3K9ac enrichment within both gene bodies and SIENAs showed already a substantial increase after 1 h. These observations suggest that reprogramming of the H3K9ac landscape might slightly precede transcriptional activation (Fig. 2C, D). Together, the data indicates that SIENA domain formation is tightly associated with the coordinated activation of clustered defense genes and may facilitate their rapid transcriptional induction following JA signaling.

### SIENAs represent broader hyperacetylated regulatory chromatin

To test whether JA-induced histone acetylation in regulatory regions is restricted to H3K9ac, we profiled the occupancy of multiple acetylation marks on histones H3, H2B, and the histone variant H2A.Z in untreated and JA-treated (2 h JA) *Arabidopsis* Col-0 seedlings using PHILO ChIP-seq. For histone H3, we examined the classical enhancer-associated mark H3K27ac as well as H3K56ac. We also profiled the acetylation of H2B (H2BK20ac), a recently described enhancer-associated histone mark in mammals (Narita et al., 2023) as well as the acetylation of H2A.Z (H2A.Zac). Our analysis focused on SIENAs that we had previously identified (Choudhary et al., 2024). Intriguingly, all examined histone acetylation marks - including H3K9ac, H3K27ac, H3K56ac, H2BK20ac, and H2A.Zac - showed JA-induced accumulation within these regions (Fig. 3A-C). Prominent MYC2 target loci displaying substantial increases across multiple acetylation marks included *MYC2* itself and *JAO2* (*JASMONIC ACID OXIDASE 2*), both of which exhibited robust enrichment of all tested acetylation marks following JA exposure (Fig. 3A, B). Within SIENAs, MYC2 DNA-binding sites themselves were largely devoid of histone modifications; however, the surrounding chromatin displayed strong JA-induced increases in acetylation (Fig. 3D). Together, these findings indicate that JA signaling establishes a broad hyperacetylated chromatin landscape at regulatory regions encompassing multiple accessible lysine residues.

**Figure 3.**
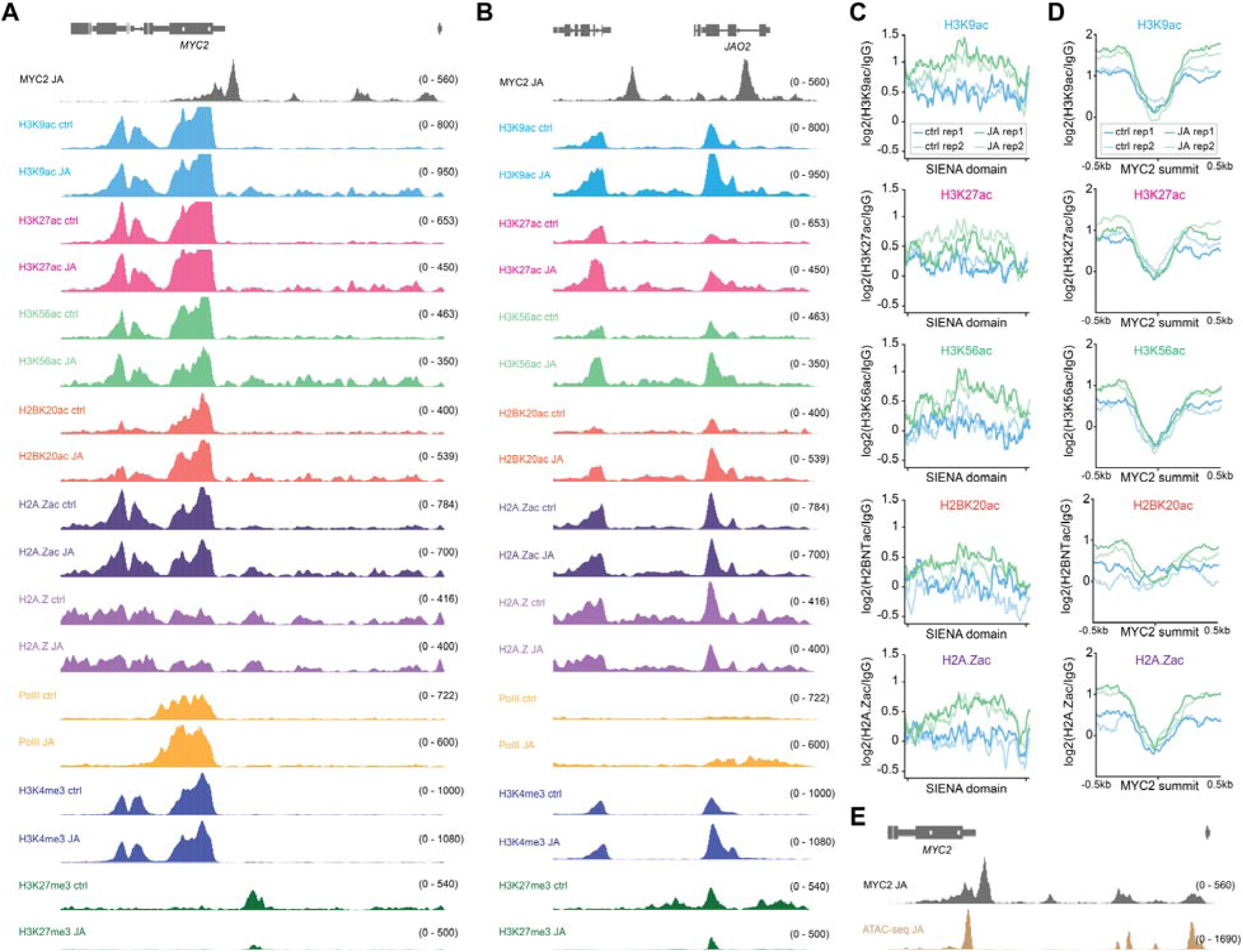
SIENAs represent broader hyperacetylated regulatory chromatin. **A, B,** Genome browser screenshot displays JA-regulated dynamics of the indicated histone marks/variants at the *MYC2* gene (**A**) as well as at the *JAO2* gene (**B**). Each track results from merging two biological replicates, and all shown tracks were normalized to their sequencing depth **C**, Metagene plots show JA-induced dynamics of indicated histone marks/variants in 30 SIENAs. Plots from two biological replicates for ctrl and 2 h JA are shown. The 30 SIENAs were scaled to 2 kb, and the respective enrichment levels were calculated as the ratio between sample and IgG control. **D,** Metagene plots show JA-induced dynamics of indicated histone marks/variants in regions spanning 500 bp both upstream and downstream of 46 MYC2 peak summits that were found in 30 SIENAs. **E,** Genome browser displays MYC2 occupancy determined with ChIP-seq and ACRs determined with ATAC-seq under inducing conditions (2h JA) at the *MYC2* gene. The ATAC-seq JA track results from merging two biological replicates. MYC2 ChIP-seq data from JA-treated *Arabidopsis* seedlings shown in **A, B**, **D**, and **E** was obtained from Choudhary et al. 2024.

In addition, we profiled the occupancies of H3K4me3, H3K27me3, H2A.Z, and RNA polymerase II (Pol II) to obtain a more comprehensive view of the JA-responsive chromatin landscape. Our analysis validated our previous findings showing that H3K4me3 and Pol II are largely absent in SIENAs and therefore display no JA-induced dynamics (Supplementary Fig. 3A). Interestingly, for H2A.Z and H3K27me3 we observed a reduction within SIENAs after JA exposure again supporting the extensive remodeling of the chromatin landscape in regulatory regions of MYC2 targets (Supplementary Fig. 3A, B).

Lastly, we performed ATAC-seq (Assay for Transposase-Accessible Chromatin using sequencing) to characterize the chromatin accessibility landscape within SIENAs. Accessible chromatin regions (ACRs) serve as a useful proxy for chromatin association of TFs and other regulatory proteins (Buenrostro et al., 2013). Because our previous analyses of the *myc234* mutant demonstrated that SIENA domain formation is largely abolished (Choudhary et al., 2024), we asked whether additional TFs might contribute to SIENA establishment. To address this question, we examined chromatin accessibility under JA conditions at 46 MYC2 binding sites located within 30 SIENAs. Remarkably, 44 of 47 ACRs (93.6%) within SIENAs overlapped with MYC2 binding sites (Supplementary Fig. 3C, D). Genome browser visualizations of the *MYC2* and *JAZ9* loci further illustrate this relationship, revealing that ACRs at JA-responsive genes coincide with MYC2 occupancy (Fig. 3E and Supplementary Fig. 3E). These findings support a model in which MYCs act as the principal architects of a highly permissive chromatin environment.

### Pharmacological disruption of proteasomal turnover compromised SIENA formation

Next, we investigated the mechanisms underlying the formation of SIENAs during active JA signaling. First, we tested whether proteasome-dependent signaling is required for SIENA domain formation. It was shown before that inhibition of the 26S proteasome with MG132 not only blocks JA-induced degradation of JAZs but also stabilizes MYC2 and increases its abundance (Zhai et al., 2013, Chung and Howe, 2009, Thines et al., 2007, Chini et al., 2007). Interestingly, MG132-stabilized MYC2 fails to fully activate target gene expression in the presence of JA indicating that MYC2 turnover is critical for its full function (Zhai et al., 2013). To test whether stabilized MYC2 can form SIENAs, we treated *Arabidopsis* Col-0 seedlings grown in liquid culture with JA (50 µM), with or without the proteasome inhibitor MG132 (50 µM) for 2 hours and subsequently profiled H3K9ac occupancies. While MG132 did not exhibit any substantial effect on the global H3K9ac landscape, JA exposure led to the expected formation of SIENAs at *MYC2* and other JA pathway components (Fig. 4A, B and Supplementary Fig. 4A, B). However, a combined JA and MG132 treatment strongly compromised SIENA domain formation (Fig. 4A, B). We still observed a modest increase in H3K9ac enrichment, which was more pronounced within the gene bodies of core MYC2 target genes (Fig. 4A, B and Supplementary Fig. 4B). This gene set was previously defined by us and comprises genes that exhibit MYC2 binding, JA-inducible transcription, and JA-induced H3K9ac accumulation within gene bodies (Choudhary et al., 2024). Our findings indicate that JA perception alone is insufficient to drive SIENA formation and that proteasome-dependent progression of the JA signaling pathway is necessary for the establishment of these hyperacetylated domains.

**Figure 4.**
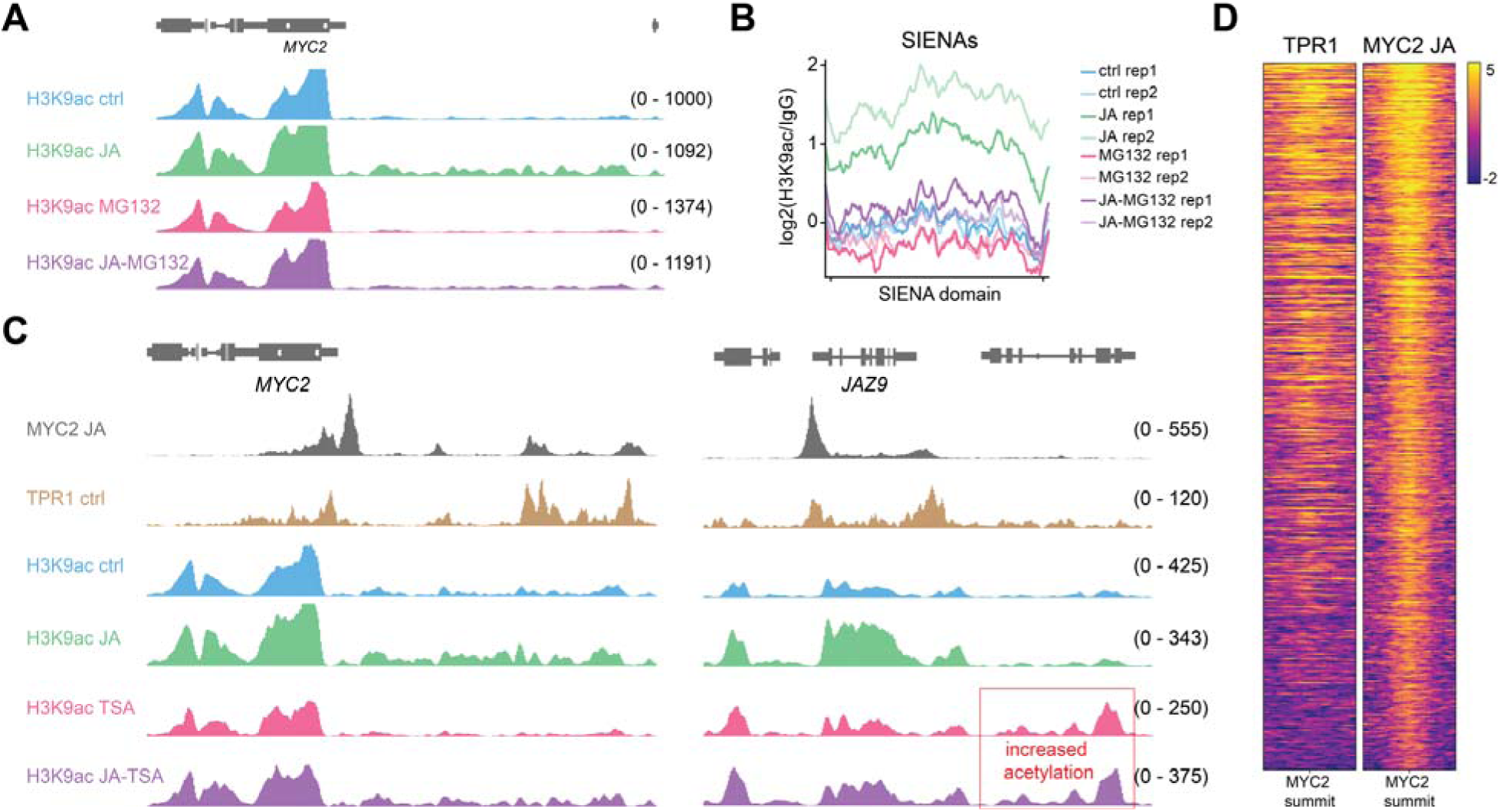
Pharmacological disruption of proteasomal turnover and HDAC activity. **A,** Genome browser shows H3K9ac dynamics at the *MYC2* gene under the indicated treatments. Ten-day-old liquid-grown *Arabidopsis* seedlings were either left untreated or treated for 2 h with 50 µM MeJA, 50 µM MG132, or a combined MeJA + MG132 treatment. Each H3K9ac ChIP-seq track was generated by merging two biological replicates, and all tracks were normalized to sequencing depth. **B**, Metagene plots show JA-induced H3K9ac dynamics under indicated treatments in 30 SIENAs. Plots from two biological replicates are shown and the 30 SIENAs were scaled to 2 kb, and the respective enrichment levels were calculated as the ratio between sample and IgG control. **C,** Genome browser shows H3K9ac dynamics at the *MYC2* (left) and the *JAZ9* (right) gene under the indicated treatments. Ten-day-old liquid-grown *Arabidopsis* seedlings were either left untreated or treated for 2 h with 50 µM MeJA, 10 µM TSA, or a combined MeJA + TSA treatment. An area showing a TSA-mediated increase of H3K9ac is also indicated. Each H3K9ac ChIP-seq track was generated by merging two biological replicates, and all tracks were normalized to sequencing depth. **D,** Heatmap shows TPR1 and MYC2 occupancy at the top 500 MYC2 binding sites (Based on MACS2 comparison, MYC2 JA vs IgG). Displayed regions span 0.5 kb upstream and 0.5 kb downstream of the MYC2 peak summit. MYC2 ChIP-seq data from JA-treated *Arabidopsis* seedlings (Choudhary et al., 2024) as well as TPR1 ChIP-seq from untreated 5-6-week-old Col-0 *TPR1-HA* plants (Griebel et al., 2023) was used for genome browser visualization in Fig. 4C and for the heatmap in Fig. 4D.

### JA overcomes TSA-induced chromatin repression at SIENA genes

The role of HDACs in JA signaling is mechanistically more complex than simple histone deacetylation, as key non-histone regulators such as TPL are also subjected to reversible acetylation (An et al., 2022). To better understand the interplay of HDACs and TPL/TPR complex in the formation of SIENAs, we treated liquid culture-grown *Arabidopsis* Col-0 seedlings grown with the broad-spectrum HDAC inhibitor trichostatin A (TSA) and subsequently profiled H3K9ac dynamics using PHILO ChIP-seq. Our rationale was to test whether HDACs operate at MYC2 target genes under non-inducing conditions. Treatment with 10 μM TSA alone for 2 h had a profound effect on the H3K9ac landscape, leading to the emergence of new acetylated regions, exemplified by the neighboring gene *JAZ9*, as well as increased H3K9ac levels at pre-existing domains throughout the genome (Fig. 4C and Supplementary Fig. 4C). Intriguingly, the effect was reversed at genes associated with SIENAs. Both gene bodies and their corresponding SIENAs exhibited lower H3K9ac levels than control samples, indicating that TSA treatment promotes a shift toward a more repressive chromatin state at these loci (Fig. 4C and Supplementary Fig. 4D, E). However, when TSA was combined with JA, SIENAs still formed to an almost similar extent as in JA-treated samples (Fig. 4C and Supplementary Fig. 4D). Despite this, H3K9ac enrichment within gene bodies remained substantially lower in JA+TSA-treated plants compared with JA treatment alone (Fig. 4C and Supplementary Fig. 4E). Together, these findings indicate that SIENA domain formation is an actively regulated JA-dependent process rather than a passive accumulation of histone acetylation. Furthermore, the restoration of SIENAs by JA in the presence of TSA suggests that JA can overcome TPL/TPR-mediated repression even under conditions of TPL hyperacetylation. Lastly, the incomplete restoration of gene-body H3K9ac indicates that SIENA domain formation and transcription-associated gene-body acetylation are at least partially separable processes when TSA is present.

Although we anticipated that TSA treatment would inhibit HDA6-mediated deacetylation associated with TPL/TPR corepressor complexes, we did not expect the pronounced shift toward a highly repressive chromatin state. Instead, we were hypothesizing that HDACs directly targeting histone proteins in SIENAs would also be inhibited thereby increasing H3K9ac levels. To further investigate the role of TPL/TPR-mediated repression at SIENA loci, we reanalyzed previously published TPR1 ChIP-seq datasets generated under non-inducing conditions and identified a high-quality dataset suitable for comparison (Griebel et al., 2023). We first examined the co-localization of MYC2 and TPR1 and although current models predict such an association, it has not previously been demonstrated on a genome-wide scale. Comparison of the top 500 MYC2 binding sites revealed a strong overlap with TPR1 occupancy, with 62.8% of MYC2 peaks co-localizing with TPR1 (Fig. 4D and Supplementary Fig. 4F, G). Focusing specifically on SIENAs, we found an even higher degree of MYC2-TPR1 co-occupancy, with 85% of MYC2 binding sites overlapping TPR1 peaks (Supplementary Fig. 4G). These findings further support a model in which HDAC-regulated TPL/TPR corepressor complexes are constitutively associated with MYC2-regulated loci prior to JA induction and actively maintain these regions in a repressed chromatin state.

## DISCUSSION

Histone acetylation is among the most extensively studied chromatin features. Yet, despite decades of research, its precise biological functions remain incompletely understood. Although numerous components of the histone acetylation machinery have been identified and partially characterized in plants, most studies have focused on histone acetylation within gene bodies, where acetylation levels are typically highest at the +1 nucleosome. Across eukaryotes, gene-body histone acetylation is strongly correlated with transcriptional activity, and in plants, transcriptional responses to environmental cues are frequently accompanied by increased histone acetylation (Willige et al., 2021, Choudhary et al., 2024, Wang et al., 2017). Whether histone acetylation is a cause or consequence of transcription remains a subject of active debate. Recent evidence supports the view that a substantial fraction of histone acetylation arises as a consequence of transcription itself. In this model, elevated histone acetylation functions as a feed-forward mechanism that reinforces and maintains high levels of gene expression (Martin et al., 2021). Consequently, it is not surprising that environmentally responsive transcriptomes are often accompanied by dynamic changes in the histone acetylation landscape.

In contrast, SIENAs represent a distinct form of histone hyperacetylation localized to regulatory regions rather than gene bodies. This distinction is solely location-based since we found that JA-induced acetylation in both regions exhibits highly similar onset kinetics, suggesting that gene body and regulatory region histone acetylation are established through a shared molecular mechanism which also seems to be a conserved feature of the chromatin landscape associated with JA-responsive genes across three plant species. Although all three tested species exhibited substantial SIENA formation, tomato displayed both the highest number of SIENAs and the largest proportion of multi-gene SIENAs. This difference cannot be simply attributed to genome size, as soybean possesses a larger genome yet contains a comparable number of single-gene SIENAs and considerably fewer multi-gene SIENAs than tomato. Interestingly, *Arabidopsis* and soybean showed a broadly similar SIENA architecture, characterized predominantly by single-gene SIENAs and only a limited number of multi-gene SIENAs. However, a notable difference among these two species was the size of individual SIENAs, particularly at key jasmonate signaling components. For example, the soybean *GmJAZ10* locus was associated with one of the largest single-gene SIENAs (~17 kb) identified in this study.

While SIENAs were initially defined based on H3K9 hyperacetylation, our comprehensive profiling of histone PTMs revealed that they are characterized by the coordinated accumulation of multiple acetylation marks, including H3K27ac, H3K56ac, H2BK20ac, and H2A.Zac. This broad acetylation signature is consistent with the generally low substrate specificity reported for many HATs. Another striking finding is the central role of MYCs in establishing SIENA domains. We previously demonstrated that H3K9ac SIENAs are dependent on MYCs (Choudhary et al., 2024) and MYC2 binding sites function as nucleation centers for SIENA formation, with acetylation spreading into the surrounding regulatory regions. Intriguingly, our ATAC-seq analyses further revealed that almost all ACRs within SIENAs overlap with MYC2 binding sites. This finding strongly suggests that the accessible chromatin landscape within SIENA domains is predominantly established and maintained by MYCs, with little evidence for substantial contributions from other TFs.

The extensive islands of histone hyperacetylation that characterize SIENAs, together with their enrichment for MYC2 binding sites, resemble classical enhancer architectures (Panigrahi and O’Malley, 2021). Recently, so-called super-enhancers (SEs) have been described in *Arabidopsis* as large genomic regions, typically exceeding 1.5 kb, that exhibit elevated chromatin accessibility (Zhao et al., 2022). However, SIENAs differ from these regions in several important aspects. First, SIENAs are highly inducible and form dynamically in response to environmental cues. Second, they are characterized by extensive histone hyperacetylation within regulatory regions, a hallmark feature of mammalian enhancers that was not reported for the previously described plant super-enhancers, including those spanning multiple genes. Thus, while SIENAs and super-enhancers share the property of encompassing extended regulatory landscapes, our findings suggest that SIENAs represent a distinct class of stimulus-responsive regulatory domains.

Two major questions remain unresolved: which HAT–HDAC modules control the dynamic establishment and removal of SIENAs, and what is the precise biological function of SIENAs. Through pharmacological inhibition of the proteasome and HDAC activity, we found that proteasome-mediated degradation of JA signaling components is critical for SIENA formation. Previous studies have shown that MYC2 is stabilized upon MG132 treatment and that MYC2 turnover is required for efficient transcriptional activation (Zhai et al., 2013). However, with our experimental setup, we cannot distinguish whether the absence of SIENAs following MG132 treatment is a direct consequence of impaired MYC2 degradation or whether inhibition of JAZ degradation more broadly prevents the JA-induced transition from a restrictive to a permissive chromatin state.

In addition, TSA treatment alone caused a pronounced reduction in H3K9ac levels at MYC2 target genes, both within gene bodies and across SIENA domains. This result was unexpected given the global increase in H3K9ac levels caused by HDAC inhibition. One possible explanation is based on the previously described role of HDA6-mediated deacetylation of TPL, which weakens the interaction between TPL and NINJA (An et al., 2022). Inhibition of HDAC activity by TSA would therefore be expected to maintain TPL in a hyperacetylated state, strengthen the TPL-NINJA interaction, and promote retention of the repressor complex at MYC2 target loci. Consistent with this model, we found that TSA treatment alone strongly reduced H3K9ac accumulation at MYC2 target genes. Notably, however, this repression could be partially overcome by JA treatment, indicating that JA-induced degradation of JAZ repressors is sufficient to alleviate repression by hyperacetylated TPL/TPR-complexes. Together, these findings support a model in which dynamic assembly and disassembly of MYC-associated repressor complexes are critical for SIENA formation.

Although the precise function of SIENAs remains unclear, it is becoming clear that SIENAs are not restricted to JA signaling. Stimulus-induced histone hyperacetylation has also been observed around PIF7 binding sites following low red to far-red (R:FR) light exposure, around HSFA1a binding sites during heat stress in tomato, and in response to drought stress in rice, suggesting that TF-governed SIENA domain formation represents a broader and potentially conserved regulatory mechanism in plants (Huang et al., 2023, Willige et al., 2021, Chang et al., 2024). Importantly, the study in rice found that promoter-associated H3K9ac can facilitate drought stress-responsive chromatin looping, functioning analogously to mammalian enhancers where histone acetylation is involved in enhancer-promoter communication through chromatin looping (Chang et al., 2024).

The sheer size of some SIENAs, the MYC dependency of SIENAs, together with the reported role of MYC2 in chromatin topology, further supports a connection between SIENA formation and dynamic 3D chromatin reorganization. Indeed, H3K4me3 ChIA-PET (Chromatin Interaction Analysis by Paired-End Tag sequencing) analyses in untreated *Arabidopsis myc2* mutants demonstrated that MYC2 contributes to higher-order chromatin organization (Deng et al., 2023). In addition, 3C (chromosome conformation capture) assays at JA-responsive loci revealed that MYC2, together with MED25, promotes chromatin looping (Wang et al., 2019). MYC2, in complex with GAME9 (GLYCOALKALOID METABOLISM 9), is also involved in facilitating communication between the distal enhancer GE1 (GAME Enhancer 1) and the GAME gene cluster, potentially through chromatin looping (Bai et al., 2024). Similarly, tomato HSFA1a has been shown to reprogram 3D chromatin architecture during heat stress, while chromatin looping has also been observed at the large regulatory region of *ATHB2*, a PIF7 target locus associated with extensive SIENA formation following low R:FR stimulation (Huang et al., 2023, Willige et al., 2021, Kim et al., 2021). Collectively, these findings point toward a role for SIENAs in higher-order chromatin organization. However, the precise mechanisms underlying their establishment, as well as their exact regulatory functions, remain largely unresolved.

## MATERIALS AND METHODS

### Genetic material and growth conditions

Seeds were surface-sterilized with bleach, rinsed in sterile water and stratified at 4 °C in the dark for 2 d. Sterilized seeds were sown on agar plates containing Linsmaier and Skoog (LS) medium (Phytotech Labs) supplemented with 1% (w/v) sucrose (Sigma-Aldrich) and grown in a Percival AR-41L3 chamber under a 16 h light / 8 h dark cycle at 19 °C. Ten-day-old seedlings were treated with gaseous methyl jasmonate (MeJA; ≥95% purity; Sigma-Aldrich) by applying 1 µl MeJA per liter of container volume onto Whatman filter paper inside a sealed transparent container; agar-plate lids were removed prior to treatment. Tissue was harvested at the indicated times and immediately processed for crosslinking. Two-week-old tomato (*Solanum lycopersicum* cv. Micro-Tom) and soybean (*Glycine max* cv. Williams 82) seedlings were grown under a 16 h light/8 h dark photoperiod at 21 °C. For methyl jasmonate (MeJA) (Sigma-Aldrich) treatments, seedlings were exposed to MeJA gaseously by applying 1 µL MeJA per liter of container volume onto Whatman filter paper placed inside a sealed transparent container. For liquid culture experiments, *Arabidopsis thaliana* Col-0 seeds were surface-sterilized with bleach, and approximately 10 mg of seeds were inoculated into 250-mL flasks containing 50 mL of LS medium. After 10 days of growth, seedlings were treated for 2 h with 50 µM MeJA, 50 µM MG132 (APExBIO), 10 µM Trichostatin A (TSA; Sigma-Aldrich), or the indicated combinations of these compounds.

### PHILO ChIP-Seq

PHILO ChIP-seq was carried out with minor modifications (Choudhary et al., 2024). We used the following antibodies for our study: anti-H3K9ac (#39137, Active Motif), anti-H3K27me3 (#39155, Active Motif), anti-H3K27ac (#39133, Active Motif), anti-H3K4me3 (#04-475, Millipore Sigma), anti-H2A.Z (#39647, Active Motif), anti-H2BK20ac (#ab177430, Abcam), anti-H3K56ac (#4243, Cell Signaling Technology), anti-H2A.Zac (#C15410202, Hologic Diagenode), and anti-RNA polymerase II 8WG16 (#664906, Biolegend). IgG (#015-000-003, Jackson ImmunoResearch) served as a control. Immunoprecipitations (IPs) were carried out in PCR 8-tube strips and for each IP, 10 μl of Dynabeads Protein A (ThermoFisher Scientific) were used. Approximately 100 mg tissue was used and extracted chromatin was fragmented with 30 cycles of sonication (30 sec on, 30 sec off) using a Bioruptor Plus (Hologic Diagenode). Only the tomato chromatin was fragmented with MNase.

### RNA-seq

Total RNA was extracted from two-week-old tomato Micro-Tom seedlings using the RNeasy Plant Mini Kit (Qiagen), and strand-specific cDNA libraries were generated with the TruSeq Stranded mRNA Library Prep Kit (Illumina) following the manufacturer’s protocol.

### ATAC-seq

ATAC-seq was performed as previously described (Hsieh et al., 2024) with minor modifications. Briefly, approximately 8 g of fresh JA-treated Col-0 seedlings were collected and processed in 2 g batches for nuclei isolation. Tagmentation reactions were carried out using 50,000 nuclei per sample and 2.5 μL of Tagmentase (Hologic Diagenode). Libraries were generated using unique dual-index (UDI) primers from Hologic Diagenode (Set 1).

### ChIP-seq analysis

ChIP-seq sequencing reads were aligned to TAIR10 (*Arabidopsis thaliana*), SLM_r2.0.pmol (*Solanum lycopersicum* Micro-Tom) (Shirasawa and Ariizumi, 2023), Glycine_max_v4.0 (*Glycine max*) using Bowtie2 (version 2.4.1) (Langmead, 2010, Langmead and Salzberg, 2012). Data was evaluated using FRiP scores, mapping rate of uniquely aligned reads, number of peaks and consistency across replicates. For each ChIP experiment, we conducted two replicates. epic2 (version 0.0.52) was used to detect genome-wide occupancies of histone modifications or histone variants using IgG samples as a control (Stovner and Saetrom, 2019). For the identification of H3K9ac domains, both replicates were first merged and then subjected to epic2 analysis using IgG samples as a control. To identify MYC2 binding peaks and MYC2 peak summit regions from previously reported *Arabidopsis* and tomato MYC2 ChIP-seq data, MACS2 (version 2.2.7.1) was used (Feng et al., 2012). MACS2 was also used for the identification of ACRs from our ATAC-seq data. *Arabidopsis* MYC2 ChIP–seq data (2 h JA treatment, biological replicate 1) were obtained from Choudhary et al. 2024. Tomato MYC2 ChIP-seq data was previously published by Du et al. (2017); biological replicate CRD029134 was used for IGV genome browser visualizations. TPR1 ChIP-seq data (GSM4497315: TPR1-HA_aGFP_r1) was also previously published (Griebel et al., 2023). For the annotation of histone domains ChIPseeker (Version 1.28.3) was used (Yu et al., 2015). Gene ontology analysis was conducted with ShinyGO (version 0.85.1) (Ge et al., 2020). SAMtools (version 1.3.1) and BEDTools (version 2.25.0) were used to process PHILO ChIP-seq replicates (Li et al., 2009, Quinlan and Hall, 2010). Heatmaps, aggregate profiles, and correlation analyses of PHILO ChIP-seq datasets were generated using deepTools (v3.5.2) (Ramirez et al., 2014).

### Identification of SIENAs

Differential H3K9ac domains between 2 h of JA treatment and control were called with epic2. We defined SIENAs as the intergenic sub-intervals of these induced epic2-identified domains, derived by subtracting overlapping gene bodies from each domain using a custom Python tool (pandas/numpy). Gene bodies were taken from the corresponding genome annotation as the full genomic span of each gene, from its first to its last annotated feature (introns included), with alternative transcripts collapsed to a single locus and strand inherited from the annotation. Within each domain, gene bodies were excised, and the remaining intergenic segments retained as SIENAs, each bounded by the nearest edge of its flanking genes with inclusive 1-bp gaps; a domain spanning N genes thus yields up to N+1 SIENAs.

Domains were retained at log2 (fold change) > 1 and FDR < 0.05, and SIENAs shorter than 1,000 bp were discarded. Each SIENA domain was classified by the gene content of its parent domain (single-gene or multi-gene) and, where genic, by its orientation: a SIENA upstream of a flanking gene’s transcription start site was designated promoter-proximal and the reciprocal interval 3′/downstream, both assigned with respect to gene strand. The carving tool and parameters are available at https://github.com/linkangit/Carve_Sienas/.

### RNA-seq analysis

Lane-level FASTQ files were concatenated per sample prior to analysis. Raw reads were quality-filtered and adapter-trimmed with fastp (v1.3.3) using default parameters (Chen et al., 2018). A genome index was built with STAR (v2.7.11b) from the *Solanum lycopersicum* reference genome SLM_r2.0.pmol and corresponding GTF annotation, with --sjdbOverhang set to read length minus one (Dobin et al., 2013). Trimmed single-end reads were aligned to the indexed genome with STAR using default parameters and coordinate-sorted BAM output. Gene-level read counts were quantified with featureCounts (Subread v2.1.1) against the GTF annotation using exon features grouped by gene_id (Liao et al., 2014). Library strandedness was empirically determined from STAR strand-specific per-gene counts and confirmed to be reverse-stranded; featureCounts was run accordingly (-s 2). Differential expression was analyzed with DESeq2 (v1.50.2) in R (Love et al., 2014). Genes with fewer than 10 counts in at least three samples were removed prior to analysis. Size-factor normalization and dispersion estimation were performed using default DESeq2 settings, with a design formula of ~ condition and control as the reference level. Each treatment (JA 15 min, 30 min, 1 h, 2 h; WD 2 h) was contrasted against Control, and log2 fold-change estimates were shrunk with the adaptive shrinkage estimator implemented in ashr v2.2.63. Genes with a Benjamini–Hochberg adjusted *P* < 0.05 and absolute log2 fold change > 1 were considered differentially expressed. Heatmaps were produced with morpheus (Broad Institute, https://software.broadinstitute.org/morpheus/) from log2-transformed, size-factor-normalized counts that were z-scored per gene across samples; rows were clustered by Euclidean distance, and samples were ordered by condition.

### ATAC-seq analyses

Single-end reads were trimmed with fastp and aligned to the corresponding reference genome with Bowtie2 (v2.4.1) (Langmead and Salzberg, 2012). PCR duplicates were removed with Picard MarkDuplicates, low-quality and organellar alignments were filtered with SAMtools (v1.3.1) (Li et al., 2009). Coverage tracks were generated with deepTools (v3.5.2) (Ramírez et al., 2014). To identify ACRs, MACS2 (version 2.2.7.1) was used (Feng et al., 2012).

## ACKNOWLEDGEMENTS

We thank Themios Chionis, Kevin Malone, and Albin Schlab for providing excellent service to our growth chambers and greenhouses.

## CONTRIBUTIONS

M.A., L.D. and M.Z. designed the research. M.A., L.D., A.C. and M.Z. performed PHILO ChIP-seq and RNA-seq experiments. M.A., L.D., and M.Z. analysed generated PHILO ChIP-seq, RNA-seq, and ATAC-seq data and performed bioinformatics analyses. M.G. extracted RNA for gene expression analyses. M.A., L.D., J.G., and H. M. performed all liquid culture experiments in *Arabidopsis*. M.A., L.D., A.C. performed tissue collection and crosslinking of various plant tissues. M.A., L.D. and M.Z. prepared the figures and wrote the manuscript.

## SUPPLEMENTARY DATA

### SUPPLEMENTARY FIGURES

**Supplementary Figure 1.**
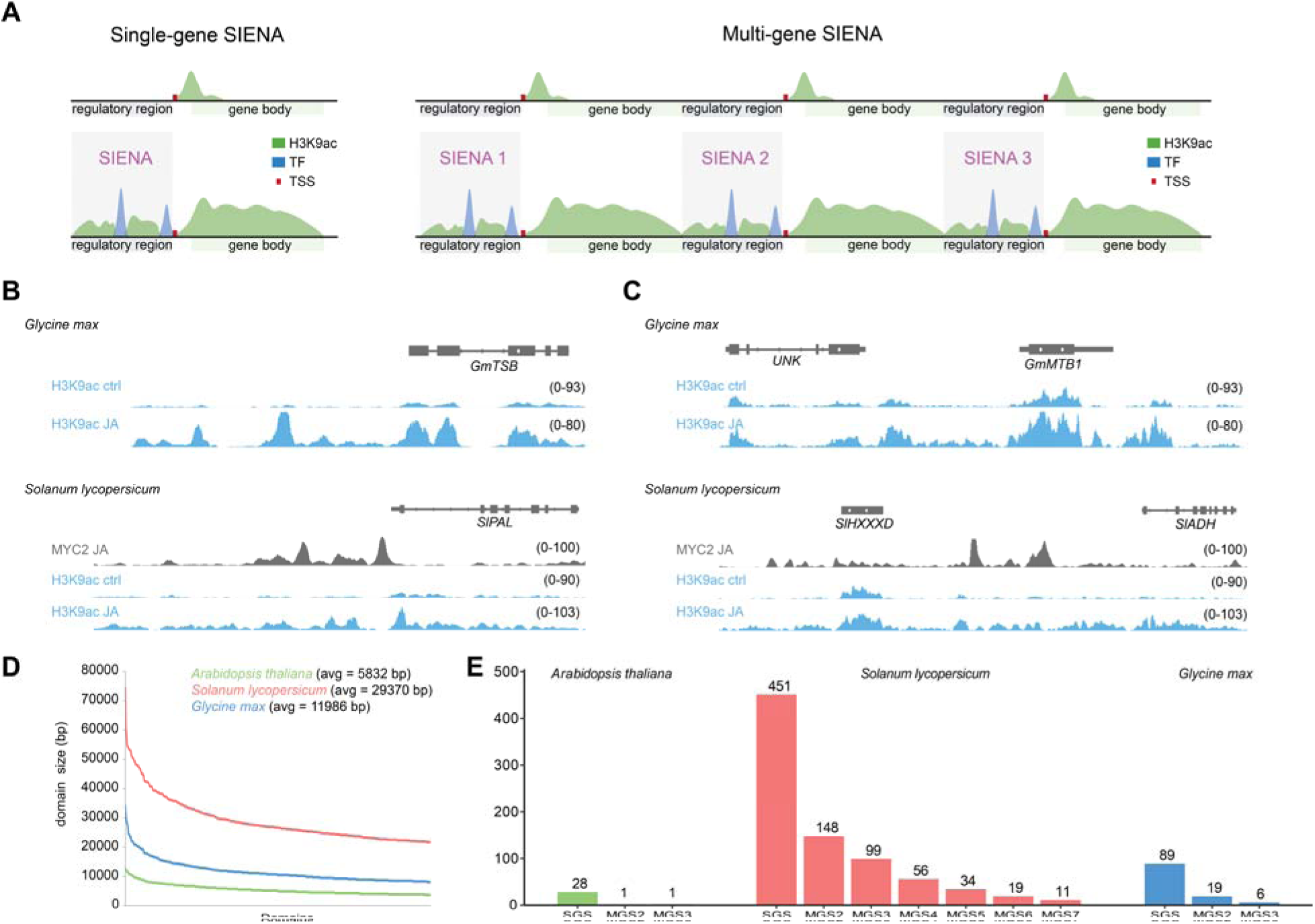
Overview of SIENA domain types. **A**, Schematic illustration of single-gene and multi-gene SIENAs. Single-gene SIENAs are defined by JA-induced H3K9ac domains that typically spread from the gene body of one gene into its regulatory regions, whereas multi-gene SIENAs comprise continuous H3K9ac domains spanning multiple adjacent genes. H3K9ac enrichment is indicated by green domains, whereas blue peaks represent TF binding peaks, which often create local valleys within SIENA domains. Illustration was generated with BioRender (https://BioRender.com/kwprjjw). **B, C,** Genome browser screenshots show examples of single-gene (**B**) and multi-gene SIENAs (**C**) in tomato and soybean. All tracks were generated by merging two biological replicates. **D,** Size distribution of the 500 largest JA-induced H3K9ac domains in *Arabidopsis*, tomato and soybean. The average (avg) domain size for each species is indicated. **E,** Bar plots show the number of single-gene and multi-gene SIENAs identified in the three indicated species. Multi-gene SIENAs are further subdivided according to the number of genes contained within each domain (MGS3 denotes a multi-gene SIENA spanning three genes).

**Supplementary Figure 2.**
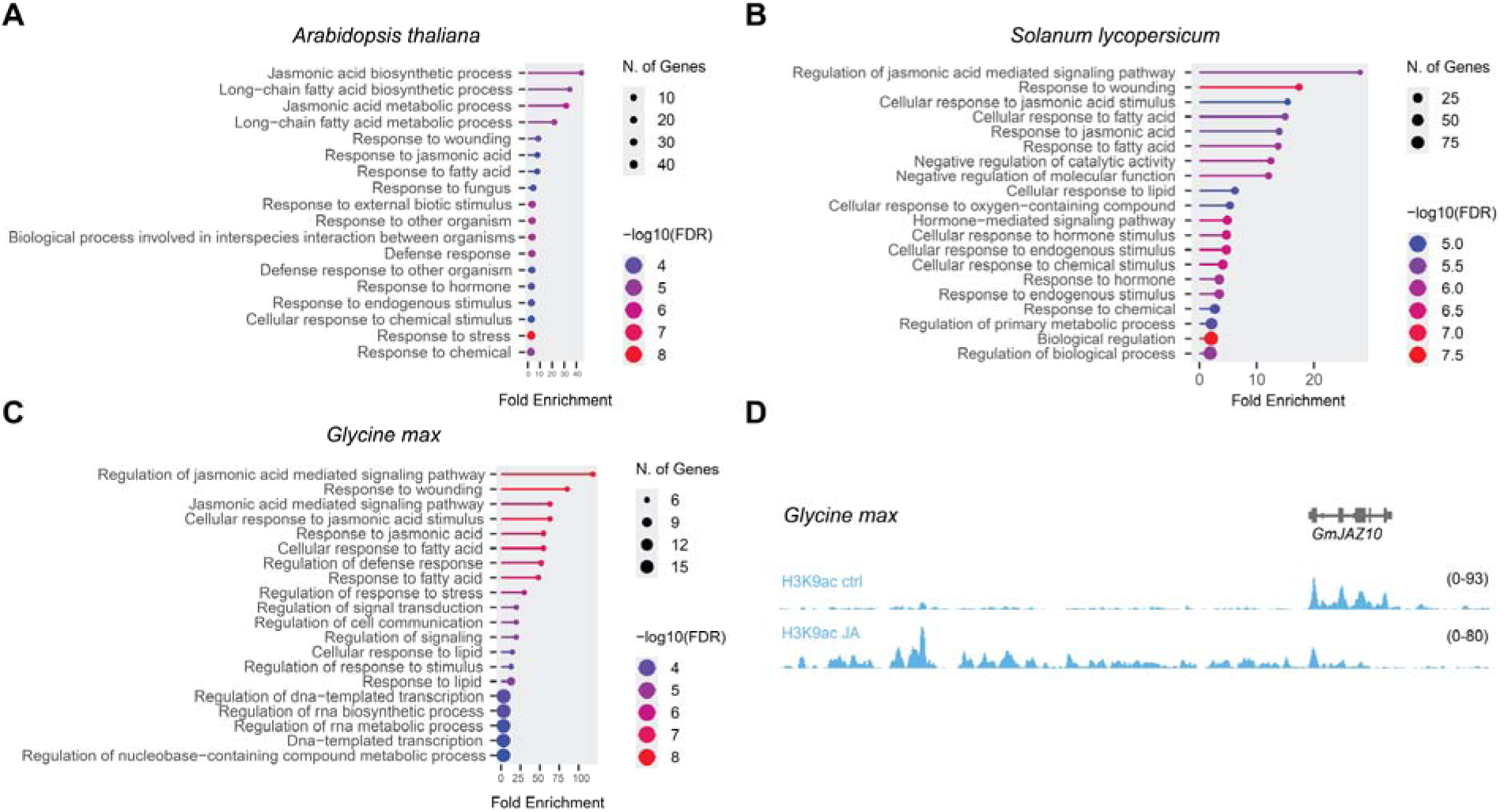
SIENAs occur at components of the JA signaling pathway. **A, B, C,** Gene ontology analysis of SIENA-associated genes in the indicated species revealed wound response and JA signaling as the predominant enriched categories. FDR values and gene counts for each category are indicated. **D,** Genome browser screenshot shows large SIENA domain at the *GmJAZ10* locus in soybean. Each track was normalized to sequencing depth and represents the merged signal from two biological replicates.

**Supplementary Figure 3.**
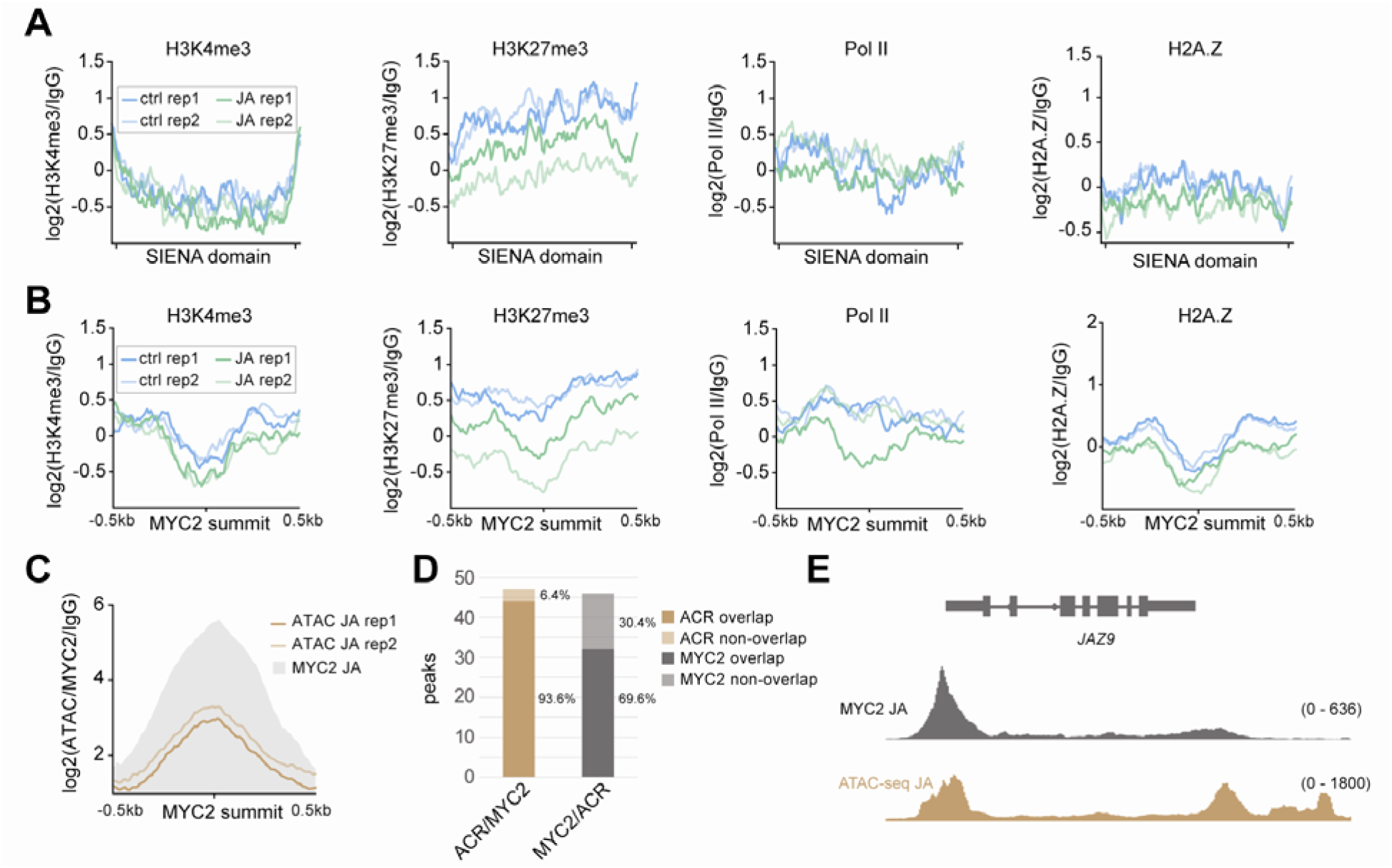
JA-induced chromatin dynamics at MYC2 target genes. **A,** Metagene plots show JA-induced dynamics of the indicated histone marks and variants across 30 *Arabidopsis* SIENAs. Profiles are shown for untreated control and 2 h JA-treated samples from two biological replicates. SIENAs were scaled to 2 kb, and enrichment levels were calculated as the ratio of each sample to the corresponding IgG control. **B,** Profiles showing JA-induced histone mark and variant dynamics across regions spanning 500 bp upstream and downstream of 46 MYC2 peak summits located within 30 SIENAs. **C,** Metagene plots showing the overlap between ACRs identified by ATAC-seq and MYC2 binding peaks across regions spanning 500 bp upstream and downstream of 46 MYC2 peak summits located within 30 SIENAs. **D,** Bar plot summarizes the overlap between ACRs and MYC2 binding sites within SIENAs. ATAC/MYC2 indicates the number of ACRs overlapping MYC2 binding sites, whereas MYC2/ATAC indicates the number of MYC2 binding sites associated with ACRs. Percentage overlap is also indicated. **E,** Genome browser view showing MYC2 occupancy determined by ChIP-seq and ACRs identified by ATAC-seq under inducing conditions at the *Arabidopsis JAZ9* locus. The JA ATAC-seq track was generated by merging two biological replicates. MYC2 ChIP-seq data was obtained from Choudhary et al. 2024.

**Supplementary Figure 4.**
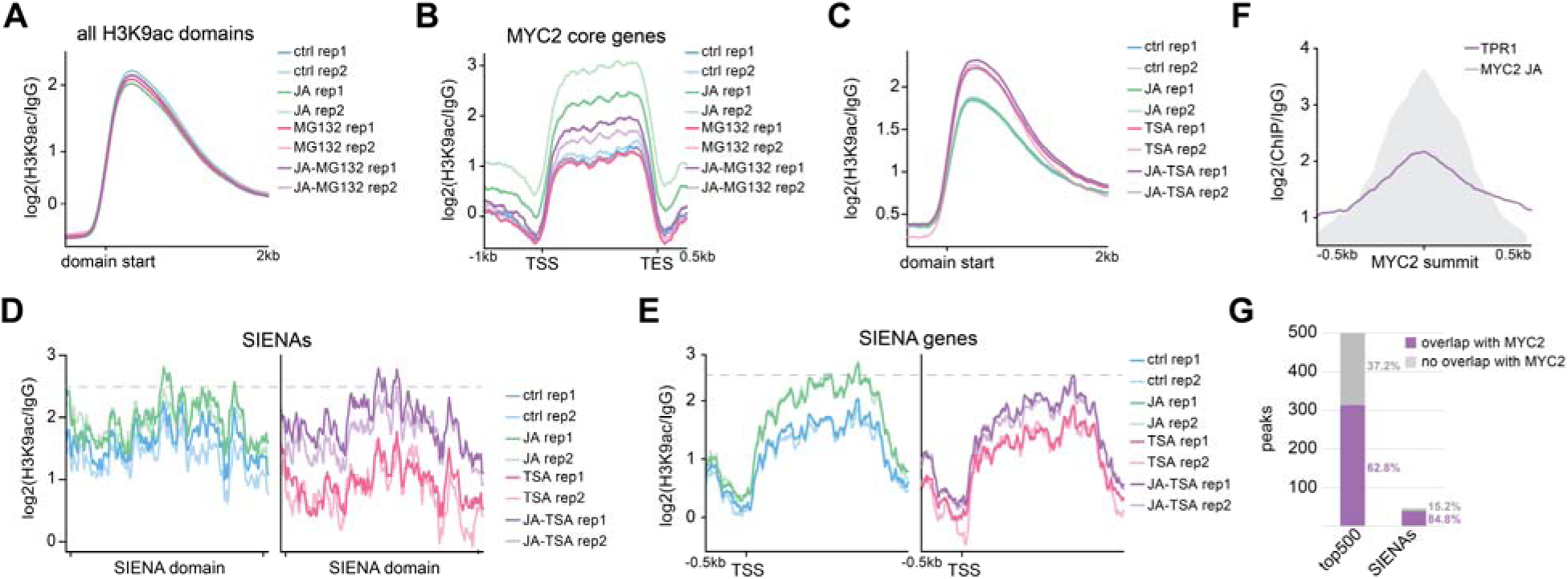
Proteasome and HDAC activity is critical for JA signaling. **A, B,** Aggregated profiles show H3K9ac occupancy across 16,452 H3K9ac domains (**A**) and 179 MYC2 core genes (**B**) under the indicated treatments (control, 2 h 50 µM MeJA, 2 h 50 µM MG132, and 2 h combined MeJA + MG132). In (**A**), regions spanning 0.5 kb upstream to 2 kb downstream of H3K9ac domain start sites are shown. In (**B**), MYC2 core genes were scaled to 2 kb, with 1 kb upstream and 0.5 kb downstream flanking regions included. **C,** Metagene plots show H3K9ac occupancy across 16,452 H3K9ac domains under the indicated treatments (control, 2 h 50 µM MeJA, 2 h 10 µM TSA, and 2 h combined MeJA + TSA). Regions spanning 0.5 kb upstream to 2 kb downstream of H3K9ac domain start sites are shown. Plots from two biological replicates are shown and the respective enrichment levels were calculated as the ratio between sample and IgG control. **D, E,** Aggregated profiles show H3K9ac occupancy across 30 SIENAs (**D**) and the corresponding gene bodies (**E**) under the indicated treatments. SIENAs and corresponding gene bodies were scaled to 2 kb. For all plots, H3K9ac ChIP-seq data from two biological replicates of untreated or treated Arabidopsis Col-0 seedlings are shown. Grey dashed lines mark approximate JA-induced H3K9ac levels. Enrichment values were calculated as the ratio between the ChIP sample and the corresponding IgG control. **F,** Metagene plots show the overlap between TPR1 and MYC2 binding across the top 500 MYC2 binding sites. Profiles are centered on the MYC2 peak summit and span 500 bp upstream and downstream. **G,** Bar plot summarizes the overlap between TPR1 and MYC2 binding sites among the top 500 MYC2 binding sites and within 46 MYC2 binding sites located across 30 *Arabidopsis* SIENAs. Percentage overlap is also indicated.

### SUPPLEMENTARY TABLES

**Supplementary Table S1 JA-responsive H3K9ac domains identified by Epic2**

**Supplementary Table S2. SIENAs derived from JA-induced H3K9ac domains in three species**

**Supplementary Table S3. Normalized expression of clustered multigene-array genes in tomato**

## REFERENCES

Ammari, M., Maseh, K. & Zander, M. 2024. PIF transcription factors-versatile plant epigenome landscapers. Frontiers in Epigenetics and Epigenomics, 2.

An, C., Deng, L., Zhai, H., You, Y., Wu, F., Zhai, Q., Goossens, A. & Li, C. 2022. Regulation of jasmonate signaling by reversible acetylation of TOPLESS in Arabidopsis. Mol Plant.

An, C., Li, L., Zhai, Q., You, Y., Deng, L., Wu, F., Chen, R., Jiang, H., Wang, H., Chen, Q. & Li, C. 2017. Mediator subunit MED25 links the jasmonate receptor to transcriptionally active chromatin. Proceedings of the National Academy of Sciences of the United States of America, 114, E8930–E8939.

Bai, F., Shu, P., Deng, H., Wu, Y., Chen, Y., Wu, M., Ma, T., Zhang, Y., Pirrello, J., Li, Z., Hong, Y., Bouzayen, M. & Liu, M. 2024. A distal enhancer guides the negative selection of toxic glycoalkaloids during tomato domestication. Nat Commun, 15, 2894.

Buenrostro, J. D., Giresi, P. G., Zaba, L. C., Chang, H. Y. & Greenleaf, W. J. 2013. Transposition of native chromatin for fast and sensitive epigenomic profiling of open chromatin, DNA-binding proteins and nucleosome position. Nature methods, 10, 1213–8.

Chang, Y., Liu, J., Guo, M., Ouyang, W., Yan, J., Xiong, L. & Li, X. 2024. Drought-responsive dynamics of H3K9ac-marked 3D chromatin interactions are integrated by OsbZIP23-associated super-enhancer-like promoter regions in rice. Genome Biol, 25, 262.

Chen, R., Jiang, H. L., Li, L., Zhai, Q. Z., Qi, L. L., Zhou, W. K., Liu, X. Q., Li, H. M., Zheng, W. G., Sun, J. Q. & Li, C. Y. 2012. The Mediator Subunit MED25 Differentially Regulates Jasmonate and Abscisic Acid Signaling through Interacting with the MYC2 and ABI5 Transcription Factors. Plant Cell, 24, 2898–2916.

Chen, S., Zhou, Y., Chen, Y. & Gu, J. 2018. fastp: an ultra-fast all-in-one FASTQ preprocessor. Bioinformatics, 34, i884–i890.

Chen, Y., Guo, P. & Dong, Z. 2024. The role of histone acetylation in transcriptional regulation and seed development. Plant Physiol, 194, 1962–1979.

Chini, A., Fonseca, S., Fernandez, G., Adie, B., Chico, J. M., Lorenzo, O., Garcia-Casado, G., Lopez-Vidriero, I., Lozano, F. M., Ponce, M. R., Micol, J. L. & Solano, R. 2007. The JAZ family of repressors is the missing link in jasmonate signalling. Nature, 448, 666–71.

Choudhary, A., Ammari, M., Yoon, H. S. & Zander, M. 2024. High-throughput capture of transcription factor-driven epigenome dynamics using PHILO ChIP-seq. Nucleic Acids Res, 52, e105.

Chung, H. S. & Howe, G. A. 2009. A critical role for the TIFY motif in repression of jasmonate signaling by a stabilized splice variant of the JASMONATE ZIM-domain protein JAZ10 in Arabidopsis. Plant Cell, 21, 131–45.

Deng, L., Zhou, Q., Zhou, J., Zhang, Q., Jia, Z., Zhu, G., Cheng, S., Cheng, L., Yin, C., Yang, C., Shen, J., Nie, J., Zhu, J. K., Li, G. & Zhao, L. 2023. 3D organization of regulatory elements for transcriptional regulation in Arabidopsis. Genome Biol, 24, 181.

Dobin, A., Davis, C. A., Schlesinger, F., Drenkow, J., Zaleski, C., Jha, S., Batut, P., Chaisson, M. & Gingeras, T. R. 2013. STAR: ultrafast universal RNA-seq aligner. Bioinformatics, 29, 15–21.

Du, M., Zhao, J., Tzeng, D. T. W., Liu, Y., Deng, L., Yang, T., Zhai, Q., Wu, F., Huang, Z., Zhou, M., Wang, Q., Chen, Q., Zhong, S., Li, C. B. & Li, C. 2017. MYC2 Orchestrates a Hierarchical Transcriptional Cascade That Regulates Jasmonate-Mediated Plant Immunity in Tomato. The Plant cell, 29, 1883–1906.

Feng, J., Liu, T., Qin, B., Zhang, Y. & Liu, X. S. 2012. Identifying ChIP-seq enrichment using MACS. Nat Protoc, 7, 1728–40.

Fernandez-Calvo, P., Chini, A., Fernandez-Barbero, G., Chico, J. M., Gimenez-Ibanez, S., Geerinck, J., Eeckhout, D., Schweizer, F., Godoy, M., Franco-Zorrilla, J. M., Pauwels, L., Witters, E., Puga, M. I., Paz-Ares, J., Goossens, A., Reymond, P., de Jaeger, G. & Solano, R. 2011. The Arabidopsis bHLH transcription factors MYC3 and MYC4 are targets of JAZ repressors and act additively with MYC2 in the activation of jasmonate responses. The Plant cell, 23, 701–15.

Ge, S. X., Jung, D. & Yao, R. 2020. ShinyGO: a graphical gene-set enrichment tool for animals and plants. Bioinformatics, 36, 2628–2629.

Griebel, T., Lapin, D., Locci, F., Kracher, B., Bautor, J., Concia, L., Benhamed, M. & Parker, J. E. 2023. Arabidopsis Topless-related 1 mitigates physiological damage and growth penalties of induced immunity. New Phytol, 239, 1404–1419.

Hickman, R., van Verk, M. C., van Dijken, A. J. H., Mendes, M. P., Vroegop-Vos, I. A., Caarls, L., Steenbergen, M., van der Nagel, I., Wesselink, G. J., Jironkin, A., Talbot, A., Rhodes, J., de Vries, M., Schuurink, R. C., Denby, K., Pieterse, C. M. J. & van Wees, S. C. M. 2017. Architecture and Dynamics of the Jasmonic Acid Gene Regulatory Network. The Plant cell, 29, 2086–2105.

Huang, Y., An, J., Sircar, S., Bergis, C., Lopes, C. D., He, X., da Costa, B., Tan, F. Q., Bazin, J., Antunez-Sanchez, J., Mammarella, M. F., Devani, R. S., Brik-Chaouche, R., Bendahmane, A., Frugier, F., Xia, C., Rothan, C., Probst, A. V., Mohamed, Z., Bergounioux, C., Delarue, M., Zhang, Y., Zheng, S., Crespi, M., Fragkostefanakis, S., Mahfouz, M. M., Ariel, F., Gutierrez-Marcos, J., Raynaud, C., Latrasse, D. & Benhamed, M. 2023. HSFA1a modulates plant heat stress responses and alters the 3D chromatin organization of enhancer-promoter interactions. Nat Commun, 14, 469.

Kim, J., Bordiya, Y., Kathare, P. K., Zhao, B., Zong, W., Huq, E. & Sung, S. 2021. Phytochrome B triggers light-dependent chromatin remodelling through the PRC2-associated PHD finger protein VIL1. Nat Plants, 7, 1213–1219.

Langmead, B. 2010. Aligning short sequencing reads with Bowtie. Current protocols in bioinformatics, Chapter 11, Unit 11 7.

Langmead, B. & Salzberg, S. L. 2012. Fast gapped-read alignment with Bowtie 2. Nature methods, 9, 357–9.

Li, H., Handsaker, B., Wysoker, A., Fennell, T., Ruan, J., Homer, N., Marth, G., Abecasis, G. & Durbin, R. 2009. The Sequence Alignment/Map format and SAMtools. Bioinformatics, 25, 2078–9.

Liao, Y., Smyth, G. K. & Shi, W. 2014. featureCounts: an efficient general purpose program for assigning sequence reads to genomic features. Bioinformatics, 30, 923–930.

Liu, Y., Du, M., Deng, L., Shen, J., Fang, M., Chen, Q., Lu, Y., Wang, Q., Li, C. & Zhai, Q. 2019. MYC2 Regulates the Termination of Jasmonate Signaling via an Autoregulatory Negative Feedback Loop. The Plant cell, 31, 106–127.

Lloyd, J. P. B. & Lister, R. 2022. Epigenome plasticity in plants. Nat Rev Genet, 23, 55–68.

Lorenzo, O., Chico, J. M., Sanchez-Serrano, J. J. & Solano, R. 2004. JASMONATE-INSENSITIVE1 encodes a MYC transcription factor essential to discriminate between different jasmonate-regulated defense responses in Arabidopsis. The Plant cell, 16, 1938–50.

Love, M. I., Huber, W. & Anders, S. 2014. Moderated estimation of fold change and dispersion for RNA-seq data with DESeq2. Genome Biol, 15, 550.

Martin, B. J. E., Brind’amour, J., Kuzmin, A., Jensen, K. N., Liu, Z. C., Lorincz, M. & Howe, L. J. 2021. Transcription shapes genome-wide histone acetylation patterns. Nat Commun, 12, 210.

Narita, T., Higashijima, Y., Kilic, S., Liebner, T., Walter, J. & Choudhary, C. 2023. Acetylation of histone H2B marks active enhancers and predicts CBP/p300 target genes. Nat Genet, 55, 679–692.

Panigrahi, A. & O’malley, B. W. 2021. Mechanisms of enhancer action: the known and the unknown. Genome Biol, 22, 108.

Pauwels, L., Barbero, G. F., Geerinck, J., Tilleman, S., Grunewald, W., Perez, A. C., Chico, J. M., Bossche, R. V., Sewell, J., Gil, E., Garcia-Casado, G., Witters, E., Inze, D., Long, J. A., de Jaeger, G., Solano, R. & Goossens, A. 2010. NINJA connects the co-repressor TOPLESS to jasmonate signalling. Nature, 464, 788–91.

Qi, T., Huang, H., Song, S. & Xie, D. 2015. Regulation of Jasmonate-Mediated Stamen Development and Seed Production by a bHLH-MYB Complex in Arabidopsis. Plant Cell, 27, 1620–33.

Quinlan, A. R. & Hall, I. M. 2010. BEDTools: a flexible suite of utilities for comparing genomic features. Bioinformatics, 26, 841–2.

Ramirez, F., Dundar, F., Diehl, S., Gruning, B. A. & Manke, T. 2014. deepTools: a flexible platform for exploring deep-sequencing data. Nucleic acids research, 42, W187–91.

Shirasawa, K. & Ariizumi, T. 2023. Near-complete genome assembly of tomato (<em>Solanum lycopersicum</em>) cultivar Micro-Tom. bioRxiv, 2023.10.26.564283.

Song, L., Huang, S. C., Wise, A., Castanon, R., Nery, J. R., Chen, H., Watanabe, M., Thomas, J., Bar-Joseph, Z. & Ecker, J. R. 2016. A transcription factor hierarchy defines an environmental stress response network. Science, 354.

Song, S. S., Huang, H., Wang, J. J., Liu, B., Qi, T. C. & Xie, D. X. 2017. is Involved in Jasmonate-Regulated Plant Growth, Leaf Senescence and Defense Responses. Plant and Cell Physiology, 58, 1752–1763.

Stovner, E. B. & Saetrom, P. 2019. epic2 efficiently finds diffuse domains in ChIP-seq data. Bioinformatics, 35, 4392–4393.

Thines, B., Katsir, L., Melotto, M., Niu, Y., Mandaokar, A., Liu, G., Nomura, K., He, S. Y., Howe, G. A. & Browse, J. 2007. JAZ repressor proteins are targets of the SCF(COI1) complex during jasmonate signalling. Nature, 448, 661–5.

Tsuda, K. & Somssich, I. E. 2015. Transcriptional networks in plant immunity. New Phytol, 206, 932–947.

Wang, H., Li, S., Li, Y., Xu, Y., Wang, Y., Zhang, R., Sun, W., Chen, Q., Wang, X. J., Li, C. & Zhao, J. 2019. MED25 connects enhancer-promoter looping and MYC2-dependent activation of jasmonate signalling. Nature plants, 5, 616–625.

Wang, L., Zhang, F., Rode, S., Chin, K. K., Ko, E. E., Kim, J., Iyer, V. R. & Qiao, H. 2017. Ethylene induces combinatorial effects of histone H3 acetylation in gene expression in Arabidopsis. BMC Genomics, 18, 538.

Willige, B. C., Zander, M., Yoo, C. Y., Phan, A., Garza, R. M., Trigg, S. A., He, Y., Nery, J. R., Chen, H., Chen, M., Ecker, J. R. & Chory, J. 2021. PHYTOCHROME-INTERACTING FACTORs trigger environmentally responsive chromatin dynamics in plants. Nat Genet, 53, 955–961.

You, Y., Zhai, Q., An, C. & Li, C. 2019. LEUNIG_HOMOLOG Mediates MYC2-Dependent Transcriptional Activation in Cooperation with the Coactivators HAC1 and MED25. Plant Cell, 31, 2187–2205.

Yu, G., Wang, L. G. & He, Q. Y. 2015. ChIPseeker: an R/Bioconductor package for ChIP peak annotation, comparison and visualization. Bioinformatics, 31, 2382–3.

Zander, M., Lewsey, M. G., Clark, N. M., Yin, L., Bartlett, A., Saldierna Guzman, J. P., Hann, E., Langford, A. E., Jow, B., Wise, A., Nery, J. R., Chen, H., Bar-Joseph, Z., Walley, J. W., Solano, R. & Ecker, J. R. 2020. Integrated multi-omics framework of the plant response to jasmonic acid. Nat Plants, 6, 290–302.

Zander, M. & Vesper, E. 2026. The environmentally responsive plant epigenome: insights from jasmonate signaling. New Phytol, 249, 2722–2728.

Zhai, Q. Z., Yan, L. H., Tan, D., Chen, R., Sun, J. Q., Gao, L. Y., Dong, M. Q., Wang, Y. C. & Li, C. Y. 2013. Phosphorylation-Coupled Proteolysis of the Transcription Factor MYC2 Is Important for Jasmonate-Signaled Plant Immunity. Plos Genetics, 9.

Zhao, H., Yang, M., Bishop, J., Teng, Y., Cao, Y., Beall, B. D., Li, S., Liu, T., Fang, Q., Fang, C., Xin, H., Nutzmann, H. W., Osbourn, A., Meng, F. & Jiang, J. 2022. Identification and functional validation of super-enhancers in Arabidopsis thaliana. Proc Natl Acad Sci U S A, 119, e2215328119.

